# Fine-scale variation in pollen availability influences *Bombus terrestris* colony behaviour, development and fitness

**DOI:** 10.1101/2025.01.28.635244

**Authors:** Harriet Lambert, Emanuel Kopp, Foteini Pashalidou, Consuelo M. De Moraes, Mark C. Mescher

**Author notes:** Corresponding author. (C.M.D.M.); (M.C.M.).

## Abstract

Climate change threatens the long-established synchronization of seasonal pollinator activity and flowering time. Bumble bees are particularly vulnerable to these shifts, with resource availability playing a significant role in colony establishment and success. Recently, we documented a mechanism whereby pollen-deprived bumble bees deliberately damaged plant leaves, speeding up the production of flowers in two different species. However, it is unknown whether bumble bees obtain a benefit through this behaviour, both on a spatial and temporal scale. In the current study, we evaluate the effects of pollen availability on colony development, behaviour and reproductive fitness in *Bombus terrestris* using semi-natural and laboratory experiments. We find that the close proximity of flowers has significant effects on reproductive investment and output of *B. terrestris* colonies in a small-scale rooftop experiment. Moreover, we find that small shifts in the timing of pollen availability (5 days) shape the behaviour and development of the first generation of offspring, with significant carry-over effects on reproductive fitness for *B. terrestris* microcolonies. This suggests that small changes in the distribution of food may have significant impacts on colony success that cannot be compensated for over the course of the season. Furthermore, our findings suggest bumble bee colonies can feasibly profit by altering the timing of pollen resources, even by a few days.

## Introduction

Climate warming and human activity rapidly change the environment, creating mismatches between plants and pollinators at spatial and temporal scales^1–5^. Studies show that a lack of flower resources reduces wild pollinator populations and pollination services locally and regionally^6–10^. These constraints particularly challenge annual pollinator species like bumble bees, which rely on timely local floral resources for colony founding and development^11–15^. Our recent discovery demonstrates that pollen-deprived bumble bee workers^16^ and founding queens^17^ damage plant leaves, accelerating flower production. This mechanism suggests direct interactions between bees and plants help maintain synchrony. This interaction could stabilize plant-pollinator communities by influencing flower production timing and seasonal availability of flower resources^18^. However, we know little about the adaptive significance of bumble bee leaf-damaging behaviour for bees or plants, or its broader ecological implications.

This study explores the fitness implications for bumble bees by investigating how small-scale changes in local resource timing affect the development, organization, and reproductive productivity of the generalist *Bombus terrestris*. As central place foragers^19^, bumble bees depend heavily on the temporal and spatial distribution of floral resources to complete their colony cycles^20^. Mated *B. terrestris* queens, like most bumble bee queens, overwinter alone and must establish new nests in early spring^21^. While damage-induced acceleration of local flower production might plausibly benefit bees later in the season by overcoming temporal gaps in flower availability, we hypothesized it is particularly relevant during nest foundation and early colony development. Pearson^22^ concludes that phenological mismatches are particularly pronounced in early spring, when flowering often precedes pollinator emergence, disrupting ecological relationships. In this resource-poor period, even small-scale shifts in flower availability can strongly affect community composition and the timing of species interactions. Moreover, research suggests that early colony development is a key predictor of success^23^ and that quantity, not quality, of resources significantly affects colony growth^24,25^.

However, the potential fitness implications of accelerated local flowering by bumblebee leaf damage remain uncertain. Work by Hemberger et al.^7^ suggests that bumble bee occurrence increases in landscapes containing more temporally continuous and abundant flowers and Bishop et al^26^. found that bumble bee abundance was positively associated with cumulative flower resources. Numerous field studies have shown that bumble bee density is reduced by flower shortages on a landscape scale^27–29^ and there is evidence that resource continuity boosts the performance of bumble bee microcolonies^30^. Moreover, Timberlake et al.^25^ conclude that bumble bees consistently forage on a diverse range of resources, even in early spring when resource is availability is low. To our knowledge, no previous study has explored how fine-scale changes in the timing or level of resource availability during the early stages of colony initiation influence subsequent development and overall success. Here, we define fine-scale changes as local resource fluctuations in close proximity to the nest (within 20 meters) and temporal variations in pollen availability over short periods, such as five-day intervals, to simulate brief environmental disruptions (e.g. a period of inclement weather).

Empirical field studies have documented strong seasonal fluctuations in food availability for pollinators^26,29,31,32^, but there is little direct evidence on the impacts of these fluctuations on bumble bee fitness. It is important to study the behavioural and reproductive responses to fluctuations in food availability, as broader biological questions of plant and pollinator species competition and coexistence may be underpinned by resource use. Additionally, measuring fitness effects and obtaining reliable estimates is a widely recognized problem in community ecology^33^. *Bombus terrestris* is highly social and polylactic species, which may select for strong behavioural flexibility, although little is known about how resource variation affects foraging choice, colony assembly, and brood production collectively.

This study aimed to address a gap in understanding the effects of fine-scale variation in resource availability on bumble bee colony development, organization, and reproduction, while assessing the potential fitness implications of accelerated flower production caused by bumble bee leaf damage. We hypothesized that small differences in the timing of resource availability during early development would have large and lasting effects on colony growth, worker behaviour, and overall reproductive success. Observing higher sensitivity to temporal variation in resource availability would support the hypothesis that bumble bee leaf-damaging behaviour is adaptive due to its effects on flowering time during colony founding.

To explore these effects, we conducted two experimental studies: (i) an outdoor experiment manipulating local flower availability for *B. terrestris* colonies, and (ii) a controlled laboratory experiment manipulating timing and abundance of pollen availability to 120 *B. terrestris* microcolonies. For a subset of these colonies, we also tracked worker behaviour. We established the resource schedule for the second experiment by constructing a statistical model estimating temporal changes in flower availability due to bumble bee leaf damage, based on *Brassica nigra* data presented in Paschalidou et al.^16^

Both experiments provide insight into the effects of floral resource availability on colony performance and fitness. The detailed observations from the large-scale laboratory study allowed us to address specific questions, including: (1) how colony foraging investment changes over time in response to food availability; (2) how task allocation among workers is influenced by constraints on pollen access; and (3) whether reproductive investment is fixed or if colonies shift the workforce into foraging rather than brood care in response to resource constraints.

## Results

### Local flower availability strongly affects colony growth and fitness

In our initial outdoor experiment, we examined how variation in flower resources near standardized colonies of founding *B. terrestris* (∼20m) influenced colony growth and fitness metrics, including the production of new reproductive gynes (females) and drones (males). We conducted these experiments on two rooftops at ETH Zurich (Switzerland): one rooftop containing garden planted with wildflowers (high flower resources) and another without flowering plants (low flowering resources), where we placed 8 establishing bumble bee colonies on each (fig. 1a). We weighed colonies twice weekly at night and examined the nest for reproductive brood cells. We also collected and measured new gynes and drones at the experiment’s endpoint, where we dissected nests to measure reproductive fitness. As predicted, local floral resources significantly affected colony growth and fitness. Colonies with access to high local flower resources had gained significantly more weight (GLMM fit by maximum likelihood, term = treatment, df = 1, Z = −2.39, P = 0.017, AIC = 2913.3) and this difference persisted throughout the experiment (fig. 1b). On June 29^th^, we removed flower resources on Roof 2 (high flower resources) by mowing the wildflower garden. Although all colonies on this roof exhibited significant weight loss after this date (GLMM fit by maximum likelihood, term = texperimental day, df = 1, Z = −3.15, P = 0.002, AIC = 2913.3), they consistently remained heavier than colonies on Roof 1(low flower resources). Flower resources also significantly affected the production of new reproductives, but this effect was only observed for gyne production which was 44% higher for colonies on Roof 2 (GLMM fit by maximum likelihood, term = treatment, df = 1, Z = −4.035, P < 0.001, AIC 345.0). Interestingly, there was a significant difference between the number of gynes and gyne cells (GLMM fit by maximum likelihood, term = treatment, df = 1, Z = −2.924, P = 0.003, AIC 345.0) see fig. 1c and fig. 1d, suggesting that colonies adopted different investment strategies for females compared to males. Colonies did not show differences in the number of drones or drone cells produced (fig. 1e and fig. 1f), indicating that gynes are more expensive to produce due to their higher feeding requirements and significant morphological dimorphism^34^. The weights of gynes and drones also varied significantly across treatments (GLMM fit by maximum likelihood, term = treatment:sex, df = 1, Z = - 3.00, P = 0.002, AIC 345.0), potentially reflecting the quality of the reproductives (see Fig. A7).

**Fig. 1.**
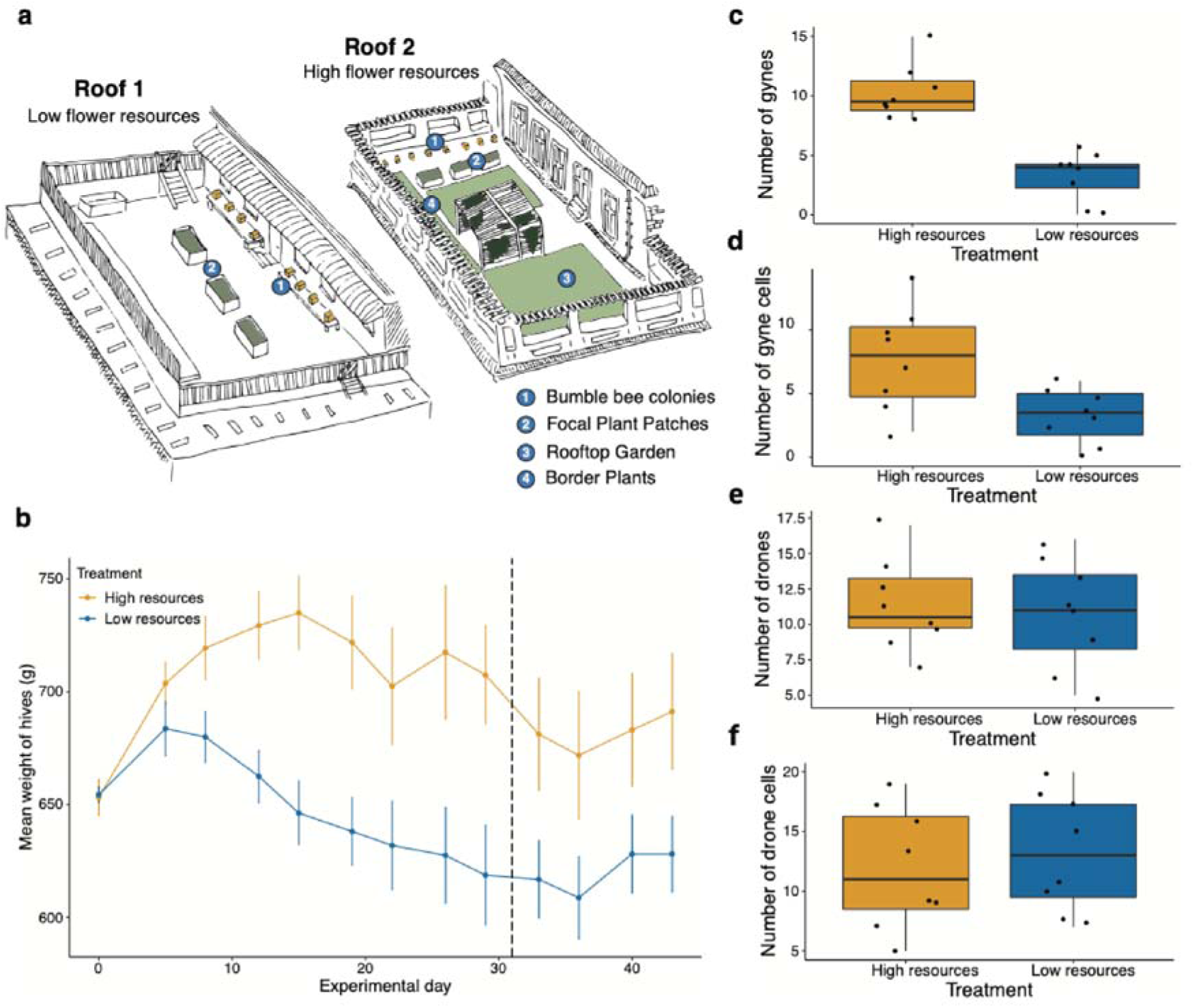
Effect of local flower availability on colony development and fitness over the course of 2019 seminatural outdoor experiment. (**a**) Experimental setup of rooftop experiment examining local proximity to flowers on colony development and fitness. (**b**) Mean weight of colonies over time. The total sample sizes for each treatment group were as follows: n=8 queenright colonies. (**c)** number of newly emerged gynes. (**d)** number of newly emerged drones. (**e**) Total number of capped gyne cells remaining in the hive at day 60 of the experiment. (**f**) Total number of capped drone cells remaining in hive at day 60 of the experiment.

Many species respond to stress by producing more of the sex with lower viability, typically favouring female production^35^. However, we found that colonies experiencing limited local flower resources shifted toward producing more males. This suggests that under resource stress, colonies bias their sex ratio toward males because producing females requires greater investment, and lower-quality colonies or queens may allocate resources to male production instead. Additionally, since the total number of reproductive brood cells did not differ significantly between treatments, this indicates a strategic adjustment in reproductive investment rather than colony failure or differential mortality. Colonies did not fail to produce females; rather, they initiated fewer gyne cells, highlighting a key reproductive strategy. The observed pattern is unlikely to result from high gyne mortality, as the significant reduction in gyne cell initiation supports an adaptive response. To further explore this strategy, we conducted laboratory studies under controlled food conditions to examine colony growth and reproductive investment in greater detail.

### Small differences in the timing of pollen availability have large effects on colony development

Our rooftop experiment demonstrates that small-scale differences in local flower availability strongly affect the development and fitness of *B. terrestris* colonies. To further explore the impact of small-scale resource variation, we conducted a high-throughput laboratory experiment using 120 *B. terrestris* microcolonies across two experimental replicates (n = 60 microcolonies per replicate). We created these microcolonies by isolating 10 bee workers in a box away from their queen, prompting one worker to become a pseudo-queen and lay unfertilized eggs, which develop into male offspring^36^. Researchers commonly use microcolonies as a model to evaluate the performance of large numbers of standardized colonies^37^, mimicking the dynamics of an establishing colony. To examine the effects of food availability, we replaced dead workers daily to maintain a consistent workforce of 10 individuals. We subjected the colonies to two treatments with different schedules for ramping up pollen resources from initially low levels. These treatment regimes, devised from a Binomial model fitted to the percentage of flowering plants per day per treatment, were conservatively estimated based on the flowering time effects of *Brassica nigra* bumblebee damage reported by Paschalidou et al. (2020). To create a conservative model based on plausible small-scale differences in flower availability, we adjusted the model fit curves to bring them closer together while maintaining similar slope shapes. The regimes included a 5-day shift in the transition from minimal to ad-libitum pollen availability (Fig. A1). We then measured worker mortality, foraging activity, offspring emergence and fitness and colony weight gain.

Our results revealed that small-scale differences in pollen availability over time had significant effects on microcolony reproduction and fitness. We first observed drone emergence on days 48 and 43 in experimental replicates 1 and 2, respectively. On average, mass hatching occurred on day 60, with a total of 256 drones hatching across both replicates. Of our initial 120 microcolonies, 53 successfully produced drones (males), including 30 in the early access treatment compared to significantly less (23) in the late access treatment. Across the total pooled microcolonies, colonies in the early access treatment produced more drones earlier than those with late access (survival curve difference test, p = 0.03, Fig. 2a). At the end of the experiment (days 67 and 70), colonies in the early access treatment had significantly more male pupae, although there was no significant difference in the number of male larvae and egg cells (Fig. 2b), indicating equal reproductive investment between the treatments. These results suggest that the timing of pollen availability is crucial for converting reproductive potential into reproductive success (Fig. A2). Given *B. terrestris* has a relatively long colony cycle compared to other bumblebee species, Ogilvie and Forrest posit that they encounter more resource turnover and gaps^38^, indicating greater sensitivity to flower shortages. We speculate that the effects of variable resource availability will have a more significant impact on queenright colonies compared to microcolonies. In queenright colonies, pollen limitation may have additive spill-over effects on colony development and offspring production, delaying the development of multiple cohorts of workers. Additionally, we observed a reduction in forewing length for drones from microcolonies in the late access treatment (one sided t-test, t = 13.229, df = 234.62, p < 0.001), along with an insignificant reduction in drone weight, see Fig. A6.

**Fig. 2.**
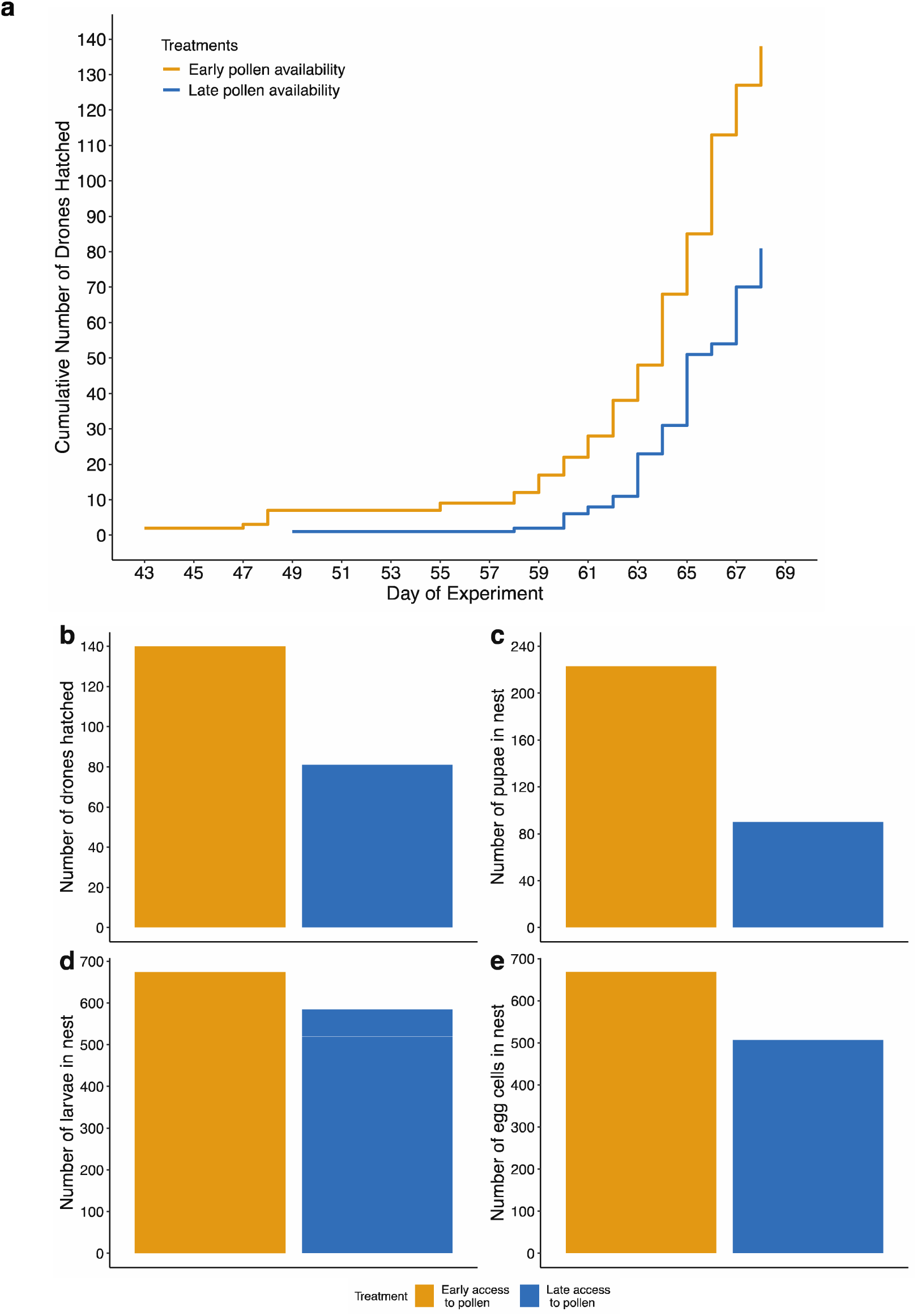
Effect of pollen availability on colony fitness and timing of reproductive emergence. (**a**) Hatching rate of drones over time according to pollen availability treatment. (**b**) Number of drones, pupae, larvae and eggs present in the nest according to pollen availability treatment.

We also observed that workers with late access to pollen had higher mortality rates than those in colonies with early access (Cox proportional hazard model, p = 0.004, SE = 0.073, Z = 2.845), and consistently lower survival probabilities (Fig. 3). Mortality refers to the occurrence of death, while survival probability represents the likelihood of an individual surviving to a specific time point. Thus, delayed pollen access reduced the likelihood of survival at any given time compared to early access. This trend persisted beyond day 25, despite all colonies receiving ad libitum pollen thereafter. Although colony weight did not significantly differ between treatments, a significant interaction between experimental day and treatment was observed (linear mixed-effects model, p = 0.019, df = 1), suggesting distinct temporal weight change patterns across treatments in both replicates (Fig. 2b). During the initial 10 days, weight gain was nearly identical; however, colonies with delayed pollen access consistently gained less weight. Even after day 50, they failed to compensate for the initial deficit and did not reach the weight of colonies with early pollen access.

**Fig. 3.**
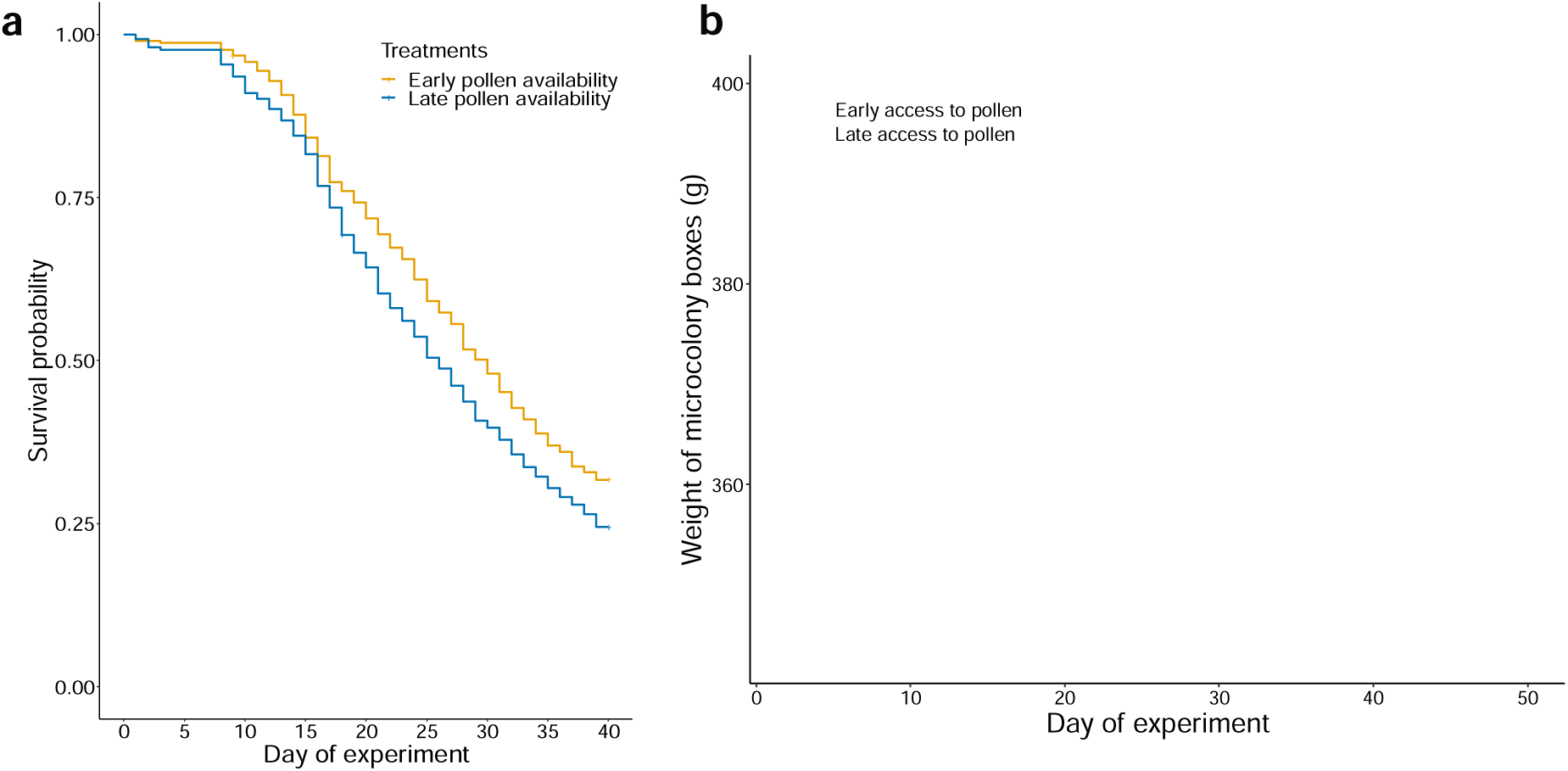
Effect of pollen availability on worker mortality and colony weight gain in microcolony experiment. (**a**) Kaplan Meier plot showing survival probability of worker individuals over time, depending on pollen availability treatment. (**b**) Effect of treatment on weight of microcolony boxes. Microcolonies in the late pollen availability treatment were not able to compensate the initial deficit and recover the weight difference despite being under *ad libitum* pollen from day 25.

### Initial differences in pollen availability have lasting effects on worker behaviour

We tracked worker behavior in 10 microcolonies (5 randomly selected from each pollen provisioning treatment in experimental replicate 1) using the image-based tracking system BEEtag^39^. We tagged 100 workers with a small QR codes and filmed the microcolonies for 10 minutes daily to monitor overall colony activity, individual foraging activity, and nursing behavior on the brood. We outlined regions of interest in the microcolony box (nectar feeder, pollen feeder, and nest) to track feeding effort. We analyzed task partitioning between days 23 and 33 of the experiment to exclude the initial phase when workers were establishing a dominance hierarchy and were extremely active (see Fig. A4).

On average, worker activity (movement distance per worker) declined over the course of the experiment as workers spent more time performing brood behaviors at the nest. However, workers from colonies with early access to pollen exhibited relatively consistent feeding effort throughout the experiment, while workers from late access colonies showed a significantly steep increase in pollen feeding effort (Fig. 4). This suggests a compensation mechanism where workers switch tasks to compensate for the initial pollen scarcity they experienced in the early period (days 1-25) by feeding more in subsequent weeks (trend towards significance in generalized mixed linear model, p = 0.0648, df =1). Colonies exhibited clear work partitioning, with a maximum of 4 workers accounting for more than 50% of the feeding effort (see Fig. A2). Worker partitioning was unaffected by treatment or worker mortality (dead workers were replaced daily to maintain a constant workforce of 10 workers), indicating that behavioral dynamics remained stable regardless of treatment. Worker partitioning in bumble bees is a crucial aspect of their social structure and efficiency, indicating that these microcolonies were functioning in an expected way.

**Fig. 4.**
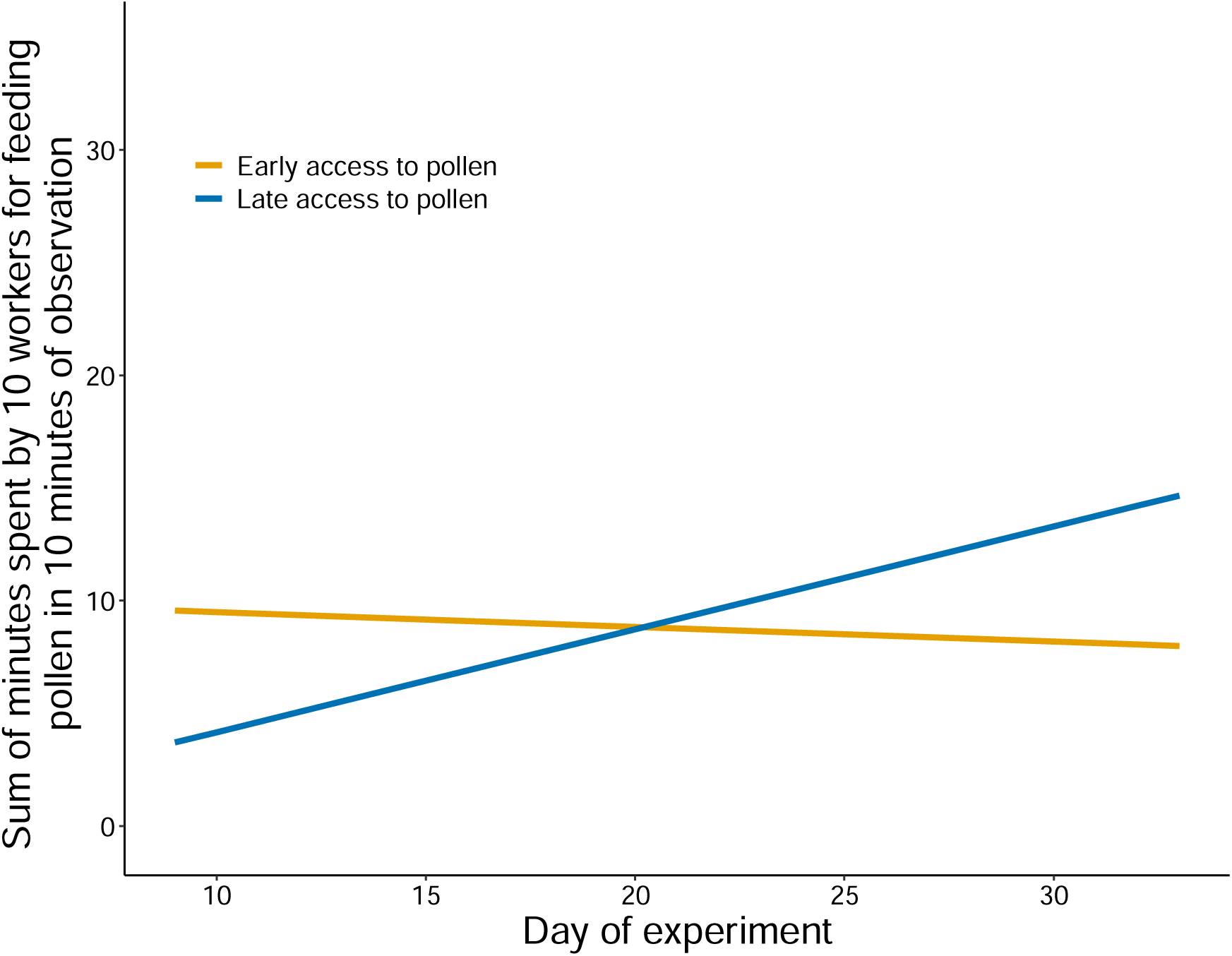
Worker activity over the course of microcolony experiment rep 1. Foraging activity remained stable for early access colonies over the course of the experiment. In contrast, late access colonies spent significantly less time near the pollen dish during the beginning of the experiment, which became offset by day 20 as pollen availability significantly increased. In the latter half of the experiment, workers from late access colonies spent more time foraging at the pollen dish (generalized linear mixed model; p = 0.065, df = 1).

## Discussion

Our research demonstrates that small-scale changes in local resource availability profoundly impact bumble bee colonies, affecting their development, organization, reproductive investment, and productivity. Our outdoor experiment revealed that very local variations in flower availability (∼20m) significantly influenced colony growth and reproductive investment. This study is among the first to suggest that differences in the onset of pollen availability, plausibly associated with leaf-damaging behaviour, have major effects on colony performance. Moreover, it indicates that if bumble bee leaf-damaging behaviour is influenced by local flower availability, colonies that can influence flowering, even on a small scale, gain a quantifiable benefit.

We found that very small-scale changes in flower availability around the nest can significantly impact colony development, sexual investment, and reproductive productivity. Surprisingly, we observed quantifiable differences on this scale, despite *B. terrestris* being a generalist capable of covering large foraging distances (although literature varies, especially for commercially reared bumble bees)^40–42^. Our colonies, placed on rooftops with 15 workers each to simulate a founding scenario, suggest that environmental disruptions or phenological mismatches in early spring, when many bumble bee species are establishing, can be particularly disruptive and have large-scale effects on colony performance. This finding adds to the body of research suggesting that resource continuity is key to reproductive success for bumble bees in the field^6,18,30,31^.

Our results indicate that rooftop colonies exhibited cohort-specific variation in sex allocation, showing remarkable flexibility in timing and production of new reproductives in response to environmental conditions. The observed difference in the number of gynes (but not drones) produced by colonies in each treatment suggests a switch in reproductive investment when colony viability is low. This counters the Trivers-Willard hypothesis, which posits that low-quality mothers would produce daughters, while high-quality mothers would produce sons^35^. Our findings suggest that bumble bees produce more males under low flower resources because males are smaller and cheaper to produce compared to gynes. This supports the idea that this is an adaptive response to resource limitation, as there was no significant difference in the number of reproductive brood cells produced between treatments. Instead, colonies initiated fewer gyne cells.

Our study provides insights into pollinator communities and their responses to climate change. The timing of pollen availability plays a crucial role in reproductive success, and our findings underscore the potential importance of synchrony between flowering times and bumble bee colony development. In the field, *Bombus terrestris* undergoes a critical early developmental period that influences colony growth^23^. However, the complex interactions between pollen access, colony behavior, and offspring fitness across the season remain largely unexplored. Although these dynamics are beginning to be disentangled^25,26^, their direct links to colony-level outcomes have yet to be explicitly established, presenting an ongoing challenge in community ecology.

Our rooftop data align with our microcolony experiments, which also show a strong dependence of colony development on fine-scale changes in resources. These experiments demonstrate that workers respond with behavioral flexibility to resource availability, shifting tasks to maximize efficiency. The foraging patterns of microcolonies indicated that colonies with early access to pollen fed consistently, while those in the late treatment showed a significant increase in feeding over time. This suggests that workers attempted to compensate for the initial pollen deficit by increasing feeding during the ad-libitum phase. However, consecutive days of unlimited pollen were inadequate to fully compensate for the previous resource shortfall. Moreover, Drones collected from microcolonies in the early access treatment had significantly longer forewings, indicating a fitness advantage for mate-seeking male bumble bees.

In conclusion, our research highlights the critical role of fine-scale resource availability in the development and success of bumble bee colonies. Even minor variations in early colony development can have profound impacts, underscoring the necessity of refining our understanding of these dynamics to bolster pollinator health and resilience. Additionally, our findings support the hypothesis that bumblebee leaf damage may help maintain synchrony between bees and plants, potentially enhancing resilience against environmental disruptions such as climate warming and habitat fragmentation.

## Methods

### Impacts of local flower availability on queenright colonies in seminatural outdoor experiment

The methodology reported for this experiment is also described in Paschalidou et al. 2020. Sixteen founding queenright *Bombus terrestris* colonies were equally divided between two treatments (flowerless and flowering) and placed on two separate rooftops (on the Zentrum campus of ETH Zurich; Zurich, Switzerland) approximately 200m from each other. Colonies on both rooftops (n = 8 per roof) were placed near focal patches of 300 flowerless plants comprising 7 different species (*Armoracia rusticana, Aurinia saxatilis, Brassica nigra, Fragaria vesca, Isatis tinctorial, Solanum lycopersicum* and *Solanum melongena)*. No plants in flower were present on Roof 1, but Roof 2 had a rooftop garden sown with mixed-species wildflowers (4.5 x 7m; ∼20m from the focal patch); in addition, we placed 30 additional flowering (i.e, currently in flower) border plants (Antirrhinum ca. 3 months old) near the focal patch on Roof 2 (Fig. S5b). All colonies on both rooftops were oriented in the same direction, with nest entrances facing West, and were sheltered from direct sunlight. Colonies were not given access to any supplemental pollen or nectar resources, except on June 17^th^ and 27^th^, when all colonies on both rooftops were provided with Biogluc sugar solution within the hive for 24h to mitigate the effects of heavy rain and hot weather (respectively). Additionally, colonies were provided with external water feeders to provide relief to the hive during hot weather. Colonies were weighed and measured twice a week (after sunset) to monitor queen presence and hive development. The rooftop garden (on Roof 2) was mowed on June 29, and we also removed the other flowering plants on Roof 2 on this date, after which point no plants in flower were present on either roof. Colonies and focal (flowerless) plant patches were kept in place until the onset of the reproductive switch point (measured as the appearance of the first drone). All hives were removed for daily monitoring in climate chambers on experimental day 45. Hives were kept at 24oC (KaClte 3000, RH 60-80%, Red lights) in small cages (50cm x 50cm x 50cm) Newly emerged reproductives were removed from the colony and frozen for measurement. At the endpoint of the experiment, all colonies were frozen and dissected to record number of egg cups, reproductive egg cups, pollen and nectar cups were present in the nest.

### Impacts of pollen availability on queenless microcolonies

#### Pollen dosage and feeding regime

Two pollen feeding regimes were devised (early pollen availability and late pollen availability) based on the flowering model described in Pashalidou et al. 2020). This model looked at the effect of *B. terrestris* worker damage on flowering time compared to mechanical damage or control plants in two species of plants (*Brassica nigra* and *solanum lycopersicum*). This model was used to simulate the effect of worker-influenced flowering, hence timing of pollen availability, under bee damaged and undamaged scenarios. However, to compensate for variation between species, the binomial model of flowering *b. nigra* was chosen as the more conservative model. Additionally, the curves of the model were modified to bring them closer together and minimise the differences between the treatments. Using this regime, microcolonies subject to late pollen availability would still reach nutrient sufficiency (*ad libitum*) by day 25, whilst colonies subject to early pollen availability would reach it by day 20. Pollen was collected from a multifloral origin (Aspermuhle Bio-BluCtenpollen, Los B0426-04), powdered on a regular basis and mixed with 30% sugar solution to form a pollen dough. This dough was weighed out each day to measure the daily dosage given to each colony (see Fig. A1).

#### Microcolony setup

For each replicate, 12 queenright maxi hives were ordered from Biobest group (Belgium) from which all workers were sourced over the course of the experiment. Each microcolony consisted of 10 workers taken from the same parent hive along with 5-7 nectar wax pots to initiate nest building. Colonies were kept at 22°C under red light conditions (KaClte 3000, RH 60%) in plastic boxes (22 cm x 15 cm x 10 cm). These boxes consisted of a nesting chamber and foraging chamber, where pollen and nectar was delivered via a plastic dish and 15 ml falcon tube that could be accessed from outside. Colonies were fed ad libitum with Biogluc® from Biobest, a sugar syrup solution (herein referred to as syrup) and refrigerated pollen dough delivered daily according to the treatment schedule. Colonies were checked daily throughout the experiment so that dead workers could be removed and replaced with new workers from the same parent hive, to ensure a consistent workforce of 10 workers per microcolony.

#### Colony monitoring and BEE-tag

In the first replicate, 5 microcolonies were selected from each treatment for BEEtag-Analysis (Crall, Gravish, et al., 2015). Each bee was tagged with a small QR-code printed on water and tear resistant paper (Xerox 003R98093). These colonies were filmed from day 1-33 using a Fujifilm XT-30 camera at highest possible resolution for 10 minutes per colony. The filming was done between 11.00 and 15.00 in accordance to diurnal colony activity levels. In each replicate, every colony was monitored daily and worker mortality recorded ( except day 39 of replicate 21). Additionally, the weight of the boxes was measured regularly over the course of the experiment. Towards the latter end of the experiment, the presence of newly emerged drones was checked daily. Offspring was easily recognized due to their silver colouration for the first twenty four hours. Drones were removed and transferred to the freezer for later measurement.

#### Dissection

All nests were dissected at the endpoint of the experiment (Day 67 for replicate 1, Day 70 for replicate 2). The number of pupae, larvae, egg cells and nectar pots were recorded before disposal. Frozen drones were taken to the laboratory and measurements on weight and length of right forewing (associated with strength of flight muscles) as a proxy for fitness.

#### Statistical analysis

All statistical analyses were carried out using R: A language and Environment for Statistical Computing version 2.15.3 (R Core Team 2013). All models were checked for autocorrelation, which was included in the model it proved to significantly increase the model fit. For Microcolony experiments, worker mortality was analysed using a Cox proportional hazard model ^34^(Therneau, 2015). The data was handled as left-truncated and right censored, where the left truncation indicates the day in which the individual starts being observed (day 1 for the 600 bumble bees at the beginning and the day in which they were used to replace a dead individual for the replacement bumble bees) and right censoring indicates the day at which the observation of the individual stopped even if the event of death didn’t happen yet (to account for the living individuals at the end of the experiment). As the individuals could not be singularly distinguished, the analysis assumed that if dead bumble bee was found in a microcolony, it was always the individual which already spent the most time in the box (i.e. the oldest individual). Outdoor experimental and Microcolony weight was analysed using linear mixed effects models (Pinheiro et al., 2020). Microcolony reproductive timing was analysed using survival curve difference tests of the G-rho family (Therneau, 2015). Differences in fitness between hatched drones were tested using one-sided t-tests.

#### BEE-tag behavioural analysis

All videos were analyzed in Matlab (MATLAB, 2019) using an adapted version of the BEE-tag script available on github (https://github.com/jamescrall/BEEtag - Crall, Gravish, et al., 2015). Videos were analyzed at a rate of 5 frames per second and the output stored as matrices of the coordinates of every tracked individual in every analyzed frame. Tracking performance was measured for each individual bumble bee by dividing the frames in which that bumble bee was tracked by the total number of analyzed frames. The first 8 days of filming were excluded from the analysis as the bumble bees were still adjusting to the microcolony setup and a pseudo-queen hierarchy was being established. Filming data of day 13 was not available and filming data for day 19 was lost due to a technical problem. In order to account for this, a gap filling approach was used in plotting, in which missing values are filled by linear interpolation between the values that bound the gap^35^ (Kelley and Richards, 2019). To compute an activity measure for each bumble bee, first the Euclidean distance between the coordinates of the same individual in subsequent frames was calculated. Then, the count of the computed distances was taken and divided by five (as the analysis was done at five frames per second) to have the number of seconds over which the total distance was tracked (one frame was assumed to last one fifth of a second). By dividing the total walked distance through the seconds over which it was recorded, the normalized distance walked in a second by every individual was obtained. As the correlation structure within the microcolonies is unknown, the results were daily averaged over every microcolony. Normalized distance was analysed using linear mixed models in R. To compute a measure for feeding effort, in every frame of every video (daily taken for each of the ten microcolonies), the number of individuals found within the defined region of interest (ROI) for pollen foraging was counted. The counts of individuals found in the ROI of all the frames (3004 per video) were summed and divided first by five to obtain the total number of seconds that the ten individuals spent in the ROI (again, one frame was assumed to last one fifth of a second) and then by 60 to obtain the same measure in minutes. This measure was called worker-minutes. Worker-minutes were analysed using linear mixed models in R.

## Acknowledgements

Many thanks to Dr. James Crall for the input regarding video tracking of bumble bees and sharing the BEEtag script with us. **Contributions:** H.L, E.K, F.G.P, M.C.M and C.M.D.M. conceived the project and designed the experiments. H.L and E.K carried out the experiments and statistical analysis. H.L and E.K carried out the modelling. All authors contributed to writing the paper. **Ethics declarations:** The authors declare no competing financial interests

## Data availability

All raw data are found on Dryad in addition to all analysis code used

## Supplementary materials for

**Fig. A1.**
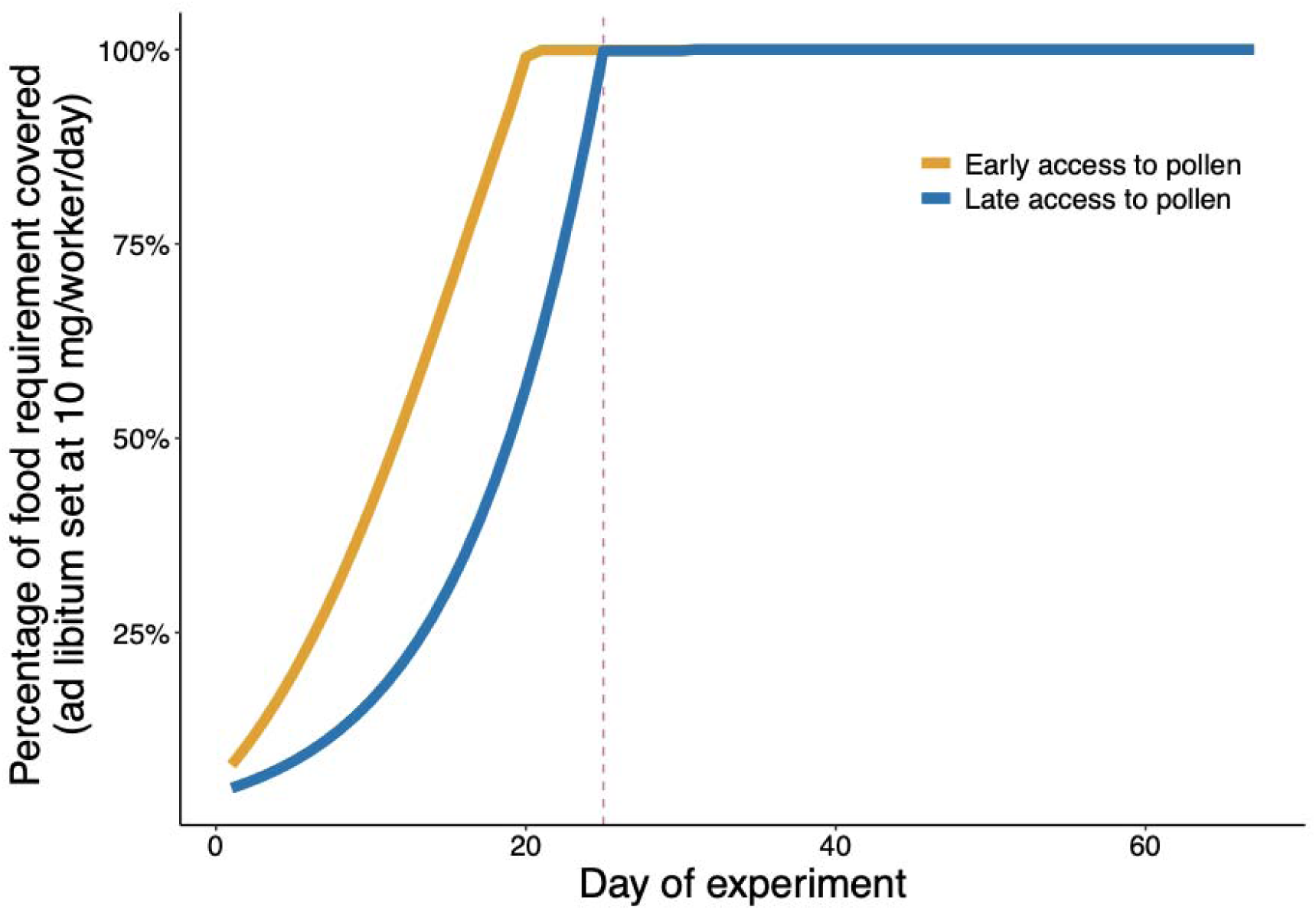
Feeding regime based on pollen availability treatments ‘Early’ and ‘Late’. Feeding regimes were created using the flowering model of *Brassica nigra* published in *Paschalidou et al. 2020*. Here, the flowering time of bee-damaged *B. nigra* plants versus undamaged control plants were compared using a GLM model with binomial error distribution and logistic-link function (*Paschalidou et al. 2020*, Fig. 1). As the flowering times between bee-damaged plants and undamaged controls varied significantly (up to 30 days difference), the feeding regime was modified to reflect a smaller difference. If treatments were chosen based on model predictions, there would have been a significant risk of falling into triviality. Here, the curves of the model fit were modified in order to minimize the differences between treatments (flatter curve than predicted by model for bee-damaged plants and steeper curve than predicted for undamaged plants), whilst maintain the shape of the curves. Moreover, for the experimental timeline (70 days), both treatments would be given pollen ad libitum for nearly two thirds of the experiment.

**Fig. A2.**
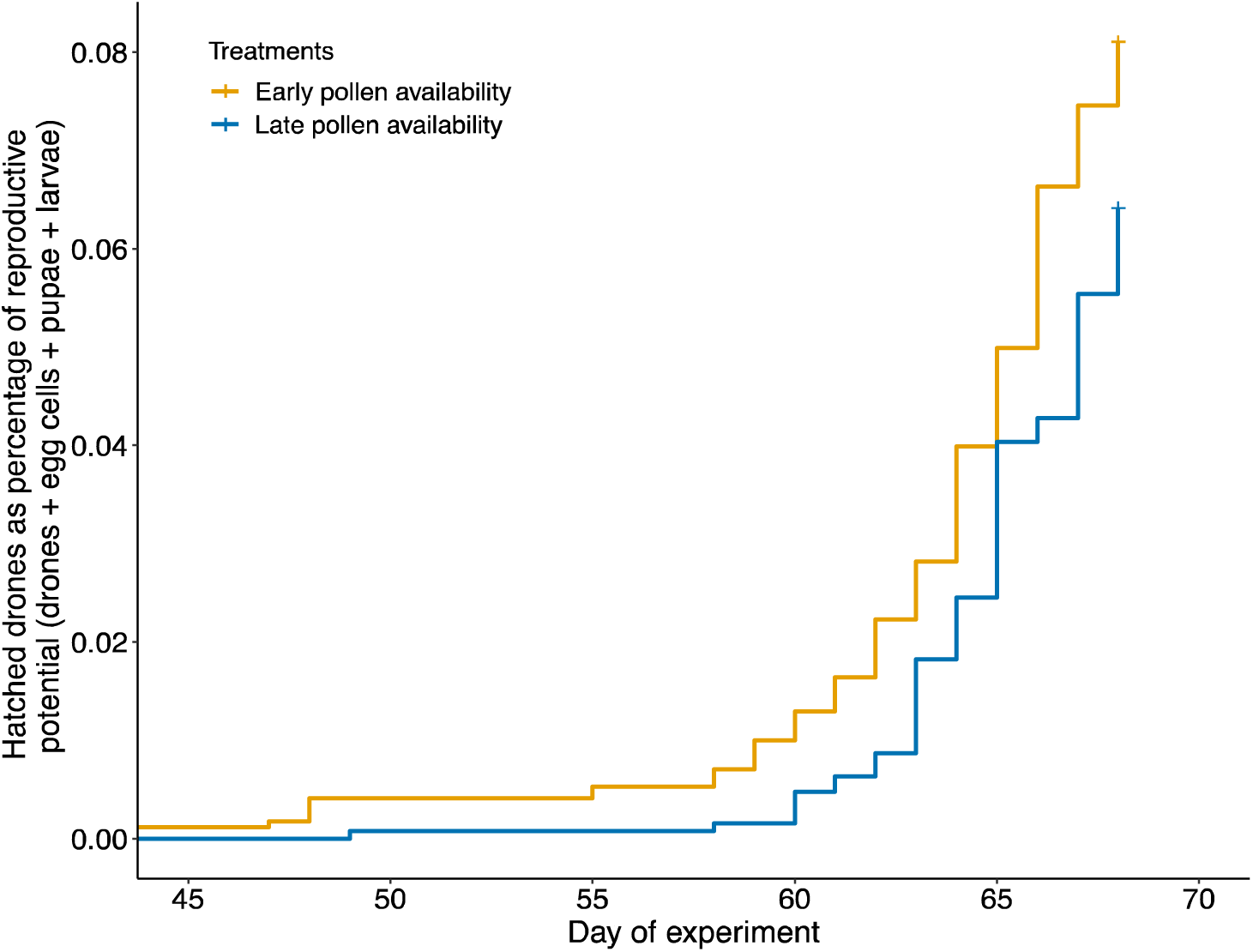
Effect of pollen availability on timing of reproductive emergence as a percentage of reproductive potential (hatched drones plus egg cells plus pupae plus larvae found in the nest at the endpoint of the experiment. Reproductive potential was plotted using the survfit function in R to generate Kaplan-Meier survival curves.

**Fig. A3.**
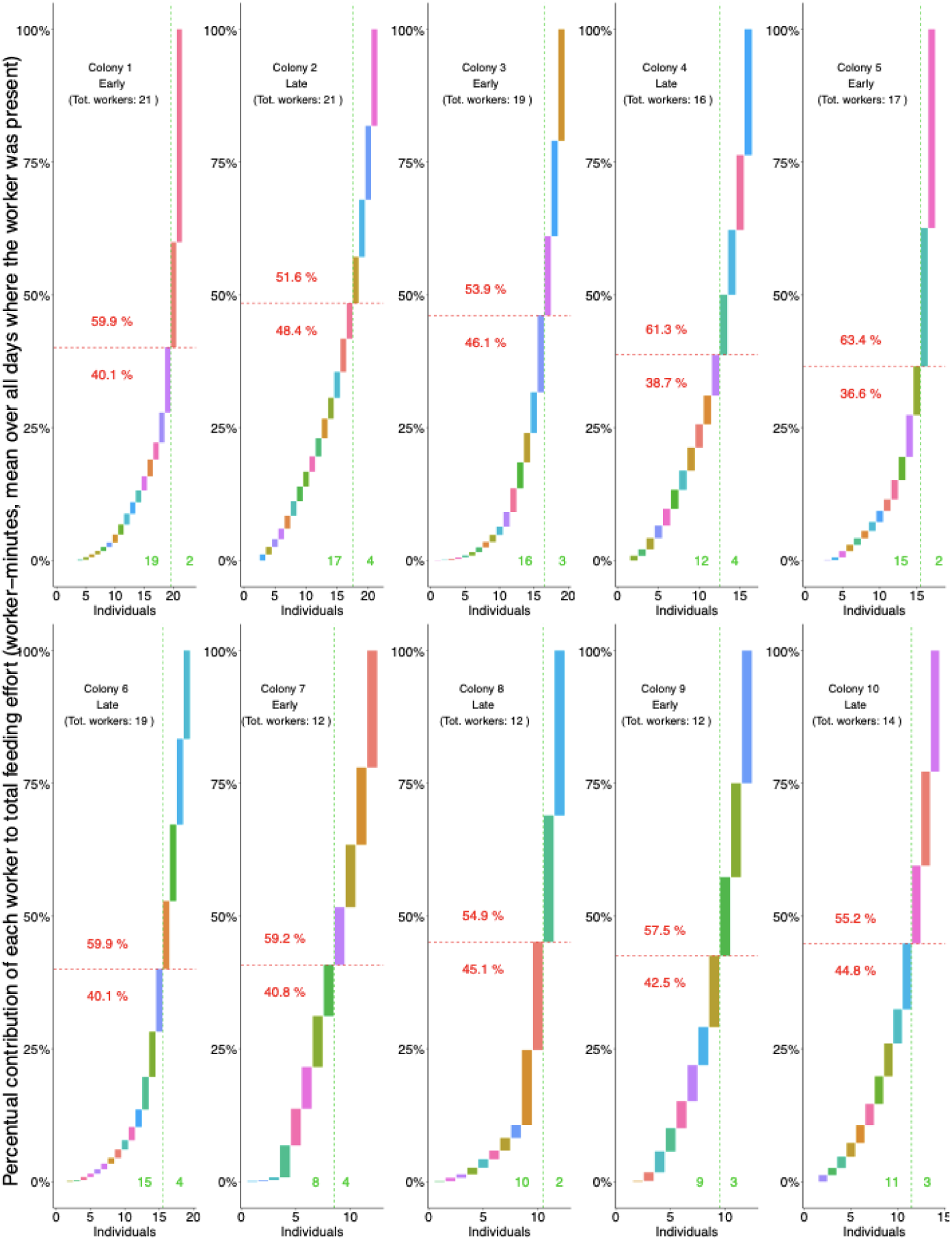
Task allocation of foraging tagged workers. Feeding behaviours were investigated in tagged workers during experimental days 23-33 of replicate 1. Feeding effort, or ‘worker-minutes’, was measured as the sum of minutes spent by every individual of the 10 worker microcolony in the pollen feeding dish, during the 10 minutes of video filmed daily. 10 randomized microcolonies were filmed overall, using 5 colonies from each pollen treatment. As the total number of workers varied between the colonies due to the differences in worker mortality, the average time per worker over the days alive was taken. A relatively high degree of task partitioning was found in microcolonies, where few workers would spend a significant period searching for pollen.

**Fig. A4.**
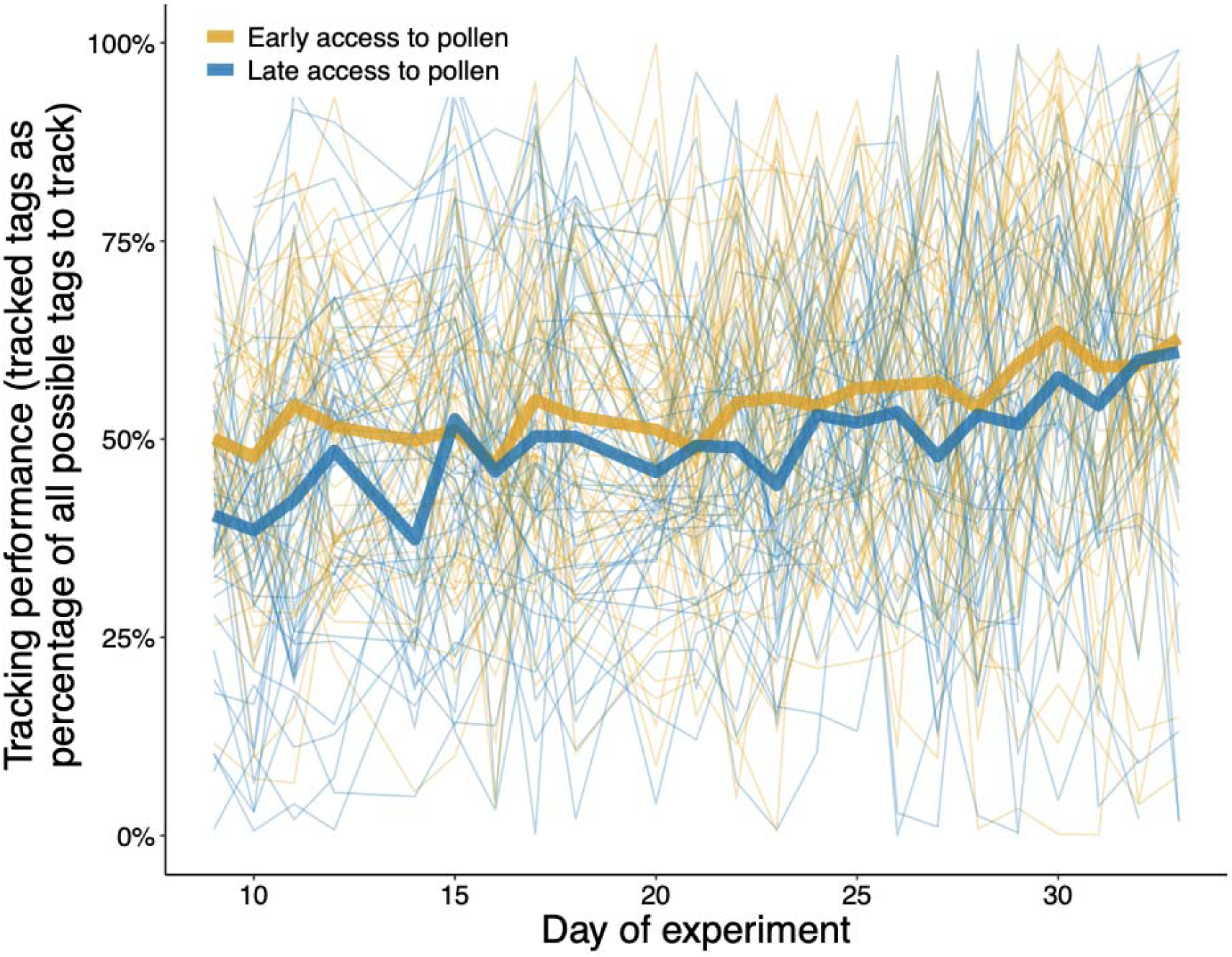
Tracking performance of bees marked using BEE-tag. Tracking was calculated daily for every worker as the % of frames in which that individual was tracked over all the frames of the video that was analysed (3004 frames per video). Overall mean tracking was ∼52%. This was influenced by a number of factors including physiological and logistical constraints. For example, individuals that had a behavioural preference for hiding under the nest were tracked less often due to the lower visibility of their tag. The trend toward increased tracking performance over the course of the experiment is presumed due to the activity changes which made tracking easier.

**Fig. A5.**
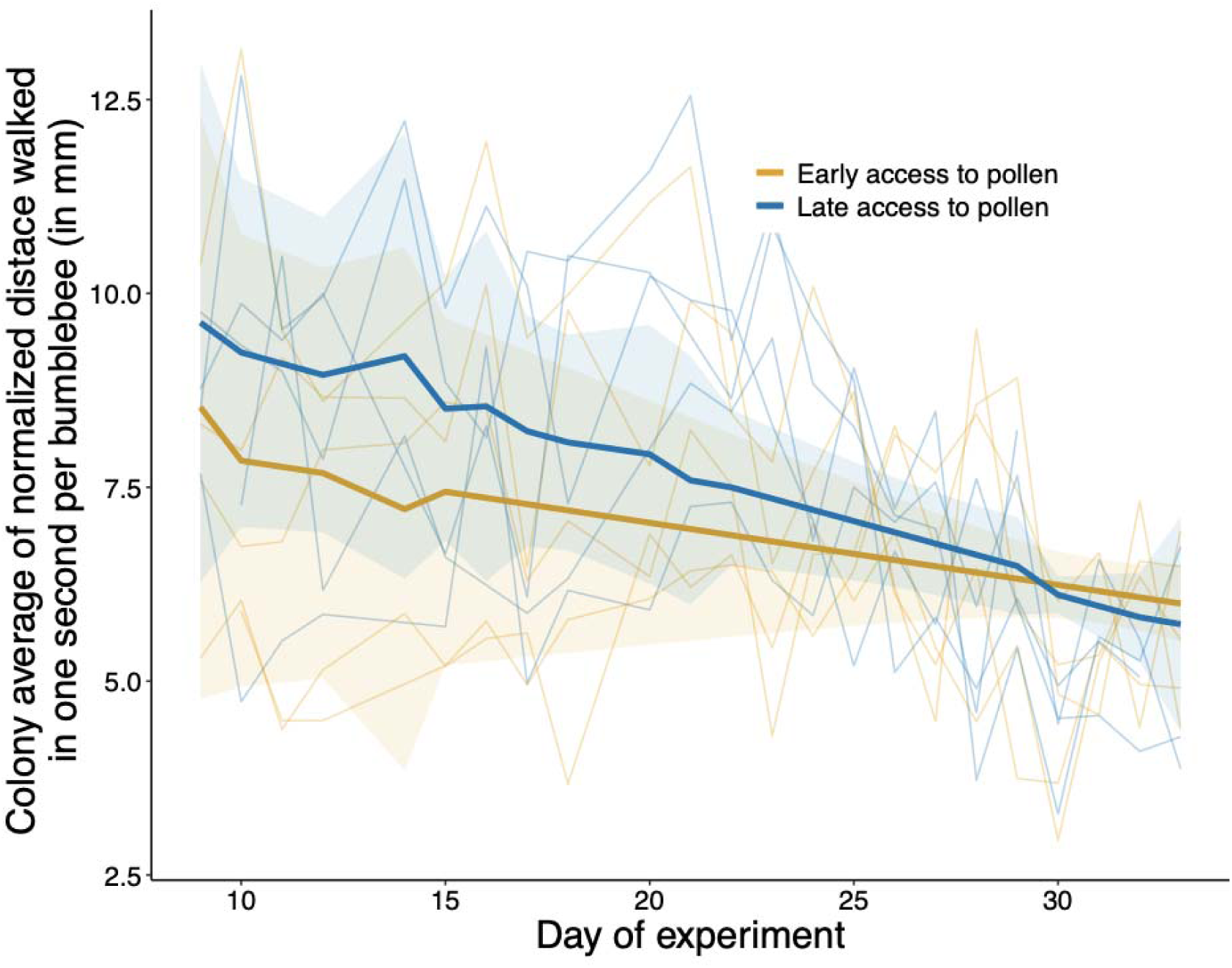
Normalized distance walked in one second per *B. terrestris* worker (colony average over the filmed experimental period. In order to measure overall colony activity, the normalized walked distance per *B. terrestris* worker was calculated and pooled per microcolony. Colonies tended to decrease in activity over the course of the experimental timeline, although this shift was stronger for colonies given the ‘Late’ pollen access treatment. This suggests they spent more time looking for food at the beginning of the experiment when experiencing pollen scarcity, which decreased when pollen became more available (ab libitum at day 25).

**Fig. A6.**
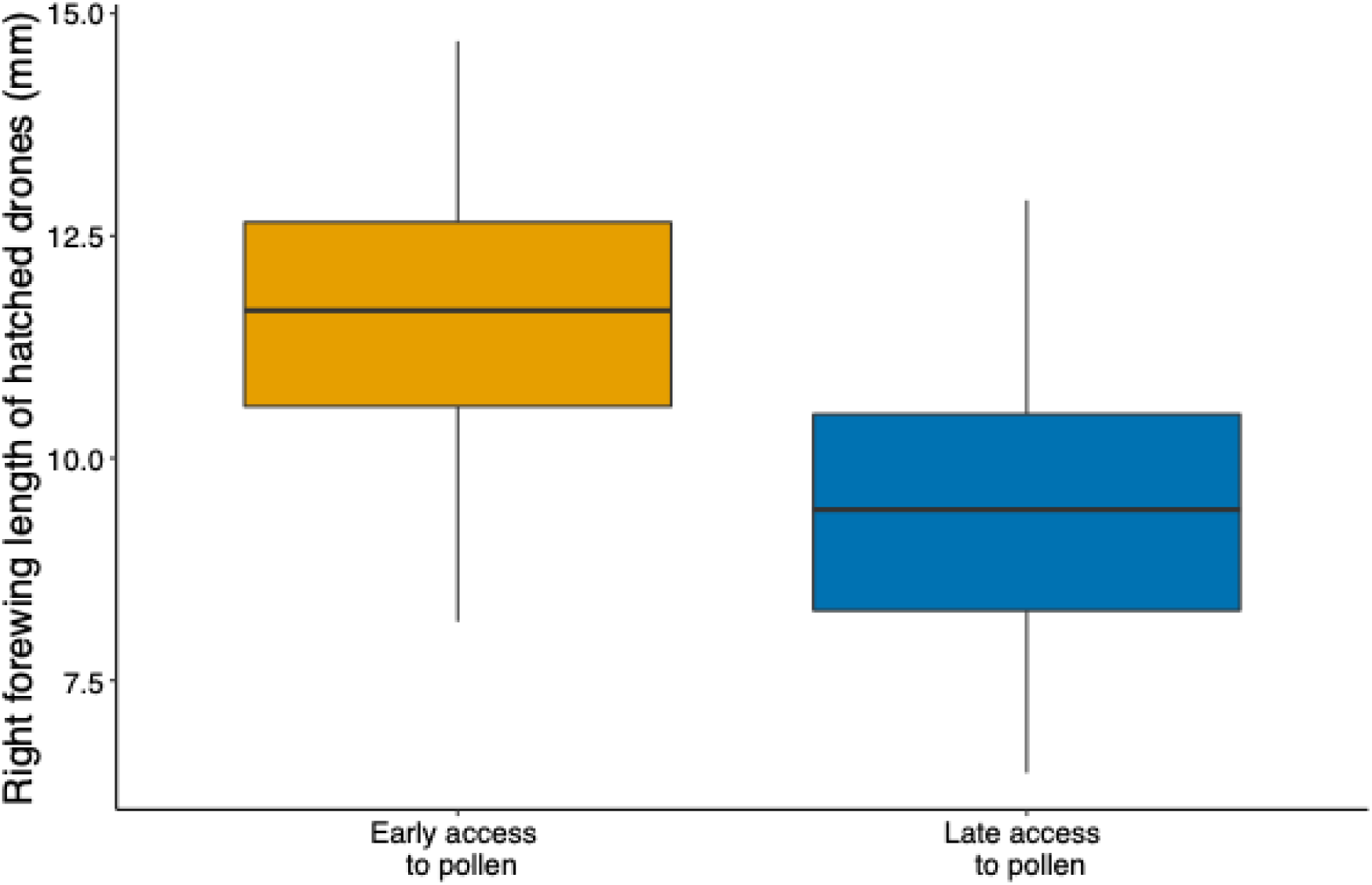
Effect of pollen Access timing on right forewinglLength of hatched drones. right forewing length (mm) of all hatched drones under two different conditions: early access to pollen and late access to pollen. The data points represent the pooled measurements of wing lengths across two experimental replicates, highlighting the impact of pollen access timing on the physical development of the drones.

**Fig. A7.**
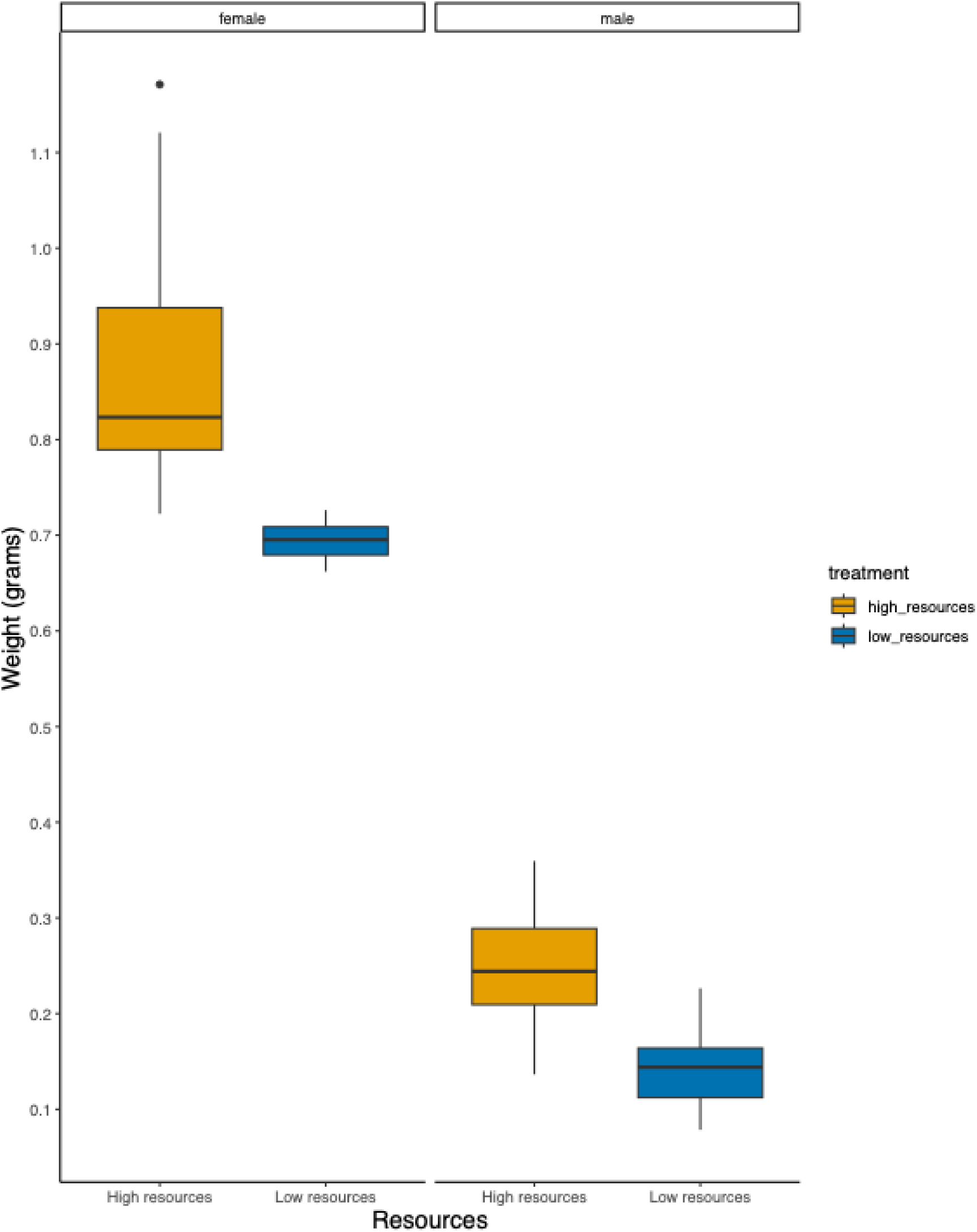
Female and male offspring weights according to access to local flower resources. Variation in offspring weights of *Bombus terrestris* colonies based on the proximity to local flower resources. The experiment was conducted on two rooftops at ETH Zurich: one with a garden of wildflowers (high flower resources) and another without flowering plants (low flower resources). We placed eight bumble bee colonies on each rooftop and monitored them twice weekly. The weights of new reproductive gynes (females) and drones (males) were measured at the experiment’s endpoint. The x-axis represents the resource treatments (high and low), while the y-axis shows the weight in grams.

## References

1. Parmesan, C. & Yohe, G. A globally coherent fingerprint of climate change impacts across natural systems. Nature 421, 37–42 (2003).

2. Potts, S. G. et al. Global pollinator declines: trends, impacts and drivers. Trends in Ecology & Evolution 25, 345–353 (2010).

3. Burkle, L. A., Marlin, J. C. & Knight, T. M. Plant-Pollinator Interactions over 120 Years: Loss of Species, Co-Occurrence, and Function. Science 339, 1611–1615 (2013).

4. Schenk, M., Krauss, J. & Holzschuh, A. Desynchronizations in bee–plant interactions cause severe fitness losses in solitary bees. Journal of Animal Ecology 87, 139–149 (2018).

5. Boggs, C. L. & Inouye, D. W. A single climate driver has direct and indirect effects on insect population dynamics. Ecology Letters 15, 502–508 (2012).

6. Hemberger, J. & Gratton, C. Floral resource discontinuity contributes to spatial mismatch between pollinator supply and pollination demand in a pollinator-dependent agricultural landscapes. Landsc Ecol (2023) doi:10.1007/s10980-023-01707-w.

7. Hemberger, J., Bernauer, O. M., Gaines-Day, H. R. & Gratton, C. Landscape-scale floral resource discontinuity decreases bumble bee occurrence and alters community composition. Ecological Applications 33, e2907 (2023).

8. Miller-Struttmann, N., Miller, Z. & Galen, C. Climate driven disruption of transitional alpine bumble bee communities. Global Change Biology 28, 6165–6179 (2022).

9. Miller-Struttmann, N. E. et al. Functional mismatch in a bumble bee pollination mutualism under climate change. Science 349, 1541–1544 (2015).

10. Gérard, M., Vanderplanck, M., Wood, T. & Michez, D. Global warming and plant– pollinator mismatches. Emerging Topics in Life Sciences 4, 77–86 (2020).

11. Kudo, G. & Ida, T. Y. Early onset of spring increases the phenological mismatch between plants and pollinators. Ecology 94, 2311–2320 (2013).

12. Scheper, J. et al. Local and landscape-level floral resources explain effects of wildflower strips on wild bees across four European countries. Journal of Applied Ecology 52, 1165– 1175 (2015).

13. Goulson, D., Nicholls, E., Botías, C. & Rotheray, E. L. Bee declines driven by combined stress from parasites, pesticides, and lack of flowers. Science 347, 1255957 (2015).

14. Kerr, J. T. et al. Climate change impacts on bumblebees converge across continents. Science 349, 177–180 (2015).

15. Memmott, J., Craze, P. G., Waser, N. M. & Price, M. V. Global warming and the disruption of plant–pollinator interactions. Ecology Letters 10, 710–717 (2007).

16. Pashalidou, F. G., Lambert, H., Peybernes, T., Mescher, M. C. & De Moraes, C. M. Bumble bees damage plant leaves and accelerate flower production when pollen is scarce. Science 368, 881–884 (2020).

17. Flury, P., Stade, S., De Moraes, C. M. & Mescher, M. C. Leaf-damaging behavior by queens and workers is widespread among bumblebee species. Communications Biology. accepted (2025).

18. Schellhorn, N. A., Gagic, V. & Bommarco, R. Time will tell: resource continuity bolsters ecosystem services. Trends in Ecology & Evolution 30, 524–530 (2015).

19. Michener, C. D. The Bees of the World. (JHU Press, 2000).

20. Moquet, L., Bacchetta, R., Laurent, E. & Jacquemart, A.-L. Spatial and temporal variations in floral resource availability affect bumblebee communities in heathlands. Biodivers Conserv 26, 687–702 (2017).

21. Rasmont, P., Ghisbain, G. & Terzo, M. Bumblebees of Europe and Neighbouring Regions. (NAP Editions, 2021).

22. Pearson, K. D. Spring- and fall-flowering species show diverging phenological responses to climate in the Southeast USA. Int J Biometeorol 63, 481–492 (2019).

23. Requier, F., Jowanowitsch, K. K., Kallnik, K. & Steffan-Dewenter, I. Limitation of complementary resources affects colony growth, foraging behavior, and reproduction in bumble bees. Ecology 101, e02946 (2020).

24. Kämper, W. et al. How landscape, pollen intake and pollen quality affect colony growth in <Emphasis Type=“Italic”>Bombus terrestris</Emphasis>. Landscape Ecol 31, 2245– 2258 (2016).

25. Timberlake, T. P., et al. Ten-a-day: Bumblebee pollen loads reveal high consistency in foraging breadth among species, sites and seasons. Ecological Solutions and Evidence 5, e12360 (2024).

26. Bishop, G. A., Fijen, T. P. M., Raemakers, I., van Kats, R. J. M. & Kleijn, D. Bees go up, flowers go down: Increased resource limitation from late spring to summer in agricultural landscapes. Journal of Applied Ecology n/a,.

27. Rundlöf, M., Persson, A. S., Smith, H. G. & Bommarco, R. Late-season mass-flowering red clover increases bumble bee queen and male densities. Biological Conservation 172, 138–145 (2014).

28. Persson, A. S. & Smith, H. G. Seasonal persistence of bumblebee populations is affected by landscape context. Agriculture, Ecosystems & Environment 165, 201–209 (2013).

29. Timberlake, T. P., Vaughan, I. P., Baude, M. & Memmott, J. Bumblebee colony density on farmland is influenced by late-summer nectar supply and garden cover. Journal of Applied Ecology n/a,.

30. Hemberger, J., Witynski, G. & Gratton, C. Floral resource continuity boosts bumble bee colony performance relative to variable floral resources. Ecological Entomology 47, 703– 712 (2022).

31. Timberlake, T. P., Vaughan, I. P. & Memmott, J. Phenology of farmland floral resources reveals seasonal gaps in nectar availability for bumblebees. Journal of Applied Ecology 56, 1585–1596 (2019).

32. Austin, A. J., Lawson-Handley, L. & Gilbert, J. D. J. Plugging the hunger gap: Organic farming supports more abundant nutritional resources for bees at critical periods. 837625 Preprint at 10.1101/837625 (2019).

33. Sponsler, D., Iverson, A. & Steffan-Dewenter, I. Pollinator competition and the structure of floral resources. Ecography 2023, e06651 (2023).

34. Beekman, M. & van Stratum, P. Bumblebee sex ratios: why do bumblebees produce so many males? Proceedings of the Royal Society of London. Series B: Biological Sciences 265, 1535–1543 (1998).

35. Trivers, R. L. & Willard, D. E. Natural Selection of Parental Ability to Vary the Sex Ratio of Offspring. Science 179, 90–92 (1973).

36. Heinrich, B. Bumblebee Economics. (Harvard University Press, 2004).

37. Klinger, E. G., Camp, A. A., Strange, J. P., Cox-Foster, D. & Lehmann, D. M. Bombus (Hymenoptera: Apidae) Microcolonies as a Tool for Biological Understanding and Pesticide Risk Assessment. Environ Entomol doi:10.1093/ee/nvz117.

38. Ogilvie, J. E. & Forrest, J. R. Interactions between bee foraging and floral resource phenology shape bee populations and communities. Current Opinion in Insect Science 21, 75–82 (2017).

39. Crall, J. D., Gravish, N., Mountcastle, A. M. & Combes, S. A. BEEtag: A Low-Cost, Image-Based Tracking System for the Study of Animal Behavior and Locomotion. PLOS ONE 10, e0136487 (2015).

40. Geib, J. C., Strange, J. P. & Galen, C. Bumble bee nest abundance, foraging distance, and host-plant reproduction: implications for management and conservation. Ecological Applications 25, 768–778 (2015).

41. Redhead, J. W. et al. Effects of habitat composition and landscape structure on worker foraging distances of five bumble bee species. Ecological Applications 26, 726–739 (2016).

42. Wolf, S. & Moritz, R. F. A. Foraging distance in Bombus terrestris L. (Hymenoptera: Apidae). Apidologie 39, 419–427 (2008).

43. Krebs, J. R. & Davies, N. B. An Introduction to Behavioural Ecology. (Blackwell Scientific Publications Ltd, 1993).

44. Benton, T. Bumblebees: The Natural History & Identification of the Species Found in Britain. (HarperCollins UK, 2006).

45. Becher, M. A., Twiston-Davies, G., Osborne, J. L. & Lander, T. A. Resource gaps pose the greatest threat for bumblebees during the colony establishment phase. Insect Conservation and Diversity n/a,.

46. Williams, N. M., Regetz, J. & Kremen, C. Landscape-scale resources promote colony growth but not reproductive performance of bumble bees. Ecology 93, 1049–1058 (2012).

